# Balancing Integration and Segregation: Structural Connectivity as a Driver of Brain Network Dynamics

**DOI:** 10.1101/2025.01.24.634823

**Authors:** Javier Palma-Espinosa, Sebastián Orellana Villouta, Carlos Coronel-Oliveros, Jean Paul Maidana, Patricio Orio

## Abstract

The brain’s ability to transition between functional states while maintaining both flexibility and stability is shaped by its structural connectivity. Understanding the relationship between brain structure and neural dynamics is a central challenge in neuroscience. Prior studies link neural dynamics to local noisy activity and mesoscale coupling mechanisms, but causal links at the whole-brain scale remain elusive. This study investigates how the balance between integration and segregation in brain networks influences their dynamical properties, focusing on multistability (switching between stable states) and metastability (transient stability over time). We analyzed a spectrum of network models, from highly segregated to highly integrated, using structural metrics like modularity, efficiency, and small-worldness. Simulating neural activity with a neural mass model and analyzing Functional Connectivity Dynamics (FCD), we found that segregated networks sustain dynamic synchronization patterns, while small-world networks, which balance local clustering and global efficiency, exhibit the richest dynamical behavior. Networks with intermediate small-worldness (*ω*) values showed peak dynamical richness, measured by variance in FCD and metastability. Using Mutual Information (MI), we quantified the structure-dynamics relationship, revealing that modularity is the strongest predictor of network dynamics, as modular architectures support transitions between dynamical states. These findings underscore the importance of the small-world architecture in brain networks, where the balance between local specialization and global integration fosters the dynamic complexity necessary for cognitive functions. By emphasizing the role of modularity, this study enhances understanding of how structural features shape neural dynamics and offers insights into disruptions linked to neurological disorders.

## Introduction

The relationship between brain structure and dynamics remains an open question in neuroscience. While it is hypothesized that structural connectivity shapes neural dynamics^1–4^, the precise nature of this interaction is still unclear. Unraveling how dynamic brain activity emerges from a relatively fixed structure is crucial for understading brain function in health and dissease. Analyzing the dynamics of simulated brain dynamics using artificial networks is an important tool for unraveling this complex relationship.

Brain connectivity, or the connectome^4^, refers to the physical links between brain regions and is most accurately represented by the Structural Connectivity (SC) matrix, derived from imaging techniques^5^. The SC matrix indicates whether two regions are connected by axonal fibers^4^. In contrast, brain activity measurements using techniques like EEG^6^ and fMRI^7^ reveal dynamic activity within each region. The relationship between these activities is often described through pairwise correlations, which uncover subnetworks of stable, synchronized activity, forming what is known as Functional Connectivity (FC).

Interesingly, relationships between subnetworks observed in SC and in FC have been found^8–10^. However, while SC is a static representation, FC is not, varying over time. As such, the FC variation is interpreted as transition between multiple states, defining a dynamical regime known as Functional Connectivity Dynamics (FCD)^11–13^. Hence, clarification on *how* the static brain structure supports the dynamical activity exhibited by the FCD, is key for understanding brain function in health and dissease.

Previous research on the causal relationship between structure and function, show that at local scale (cortical areas), the observed global dynamics may emerge from noisy activity of neurons and synapses, as well as their chaotic nature^14–18^. At mesoscale (coupling between areas), global coupling^15,19,20^ and internode delay^19,21^ could be the mechanisms that drive the system towards different dynamical states. Nevertheless, while some studies have established a link between a given SC and its associated FCD^10,21,22^, a causal relationship between the topological properties of SC, namely integration and segregation, and FCD’s dynamical characteristics, such as multistability or metastability, has not been systematically studied at global scale (whole brain).

In this study, we aimed to investigate how SC’s integration and segregation correlate with FCD’s multistability and metastability. By using a wilson-cowan neural mass model over different network topologies, including a binarized version of the human connectome, we investigated how the broad spectrum of structural segregation and integration, defined by the Small-world index *ω*^23^, shapes network dynamics. For that, we compared metrics of network structure, such as Path Length, clustering coefficient, small-world index, and modularity with parameters of network dynamics, such as metastability and variance in FCD.

## Results

### Set of Structural Connectivities

We built a set of adjacency matrices that span the axis of the integration-segregation property. This set comprises five types of networks or generating algorithms: Modular networks, Hierarchical networks, Watts-Strogatz small world networks, Barabasi-Alberts scale-free networks. Each of these network-generating algorithms can be tuned to obtain a different degree of integration or segregation in the network. In addition, a binarized Human connectome following the Schaeffer-200 parcellation was used, which was also perturbed to obtain integrated or segregated versions (Fig 1).

**Figure 1.**
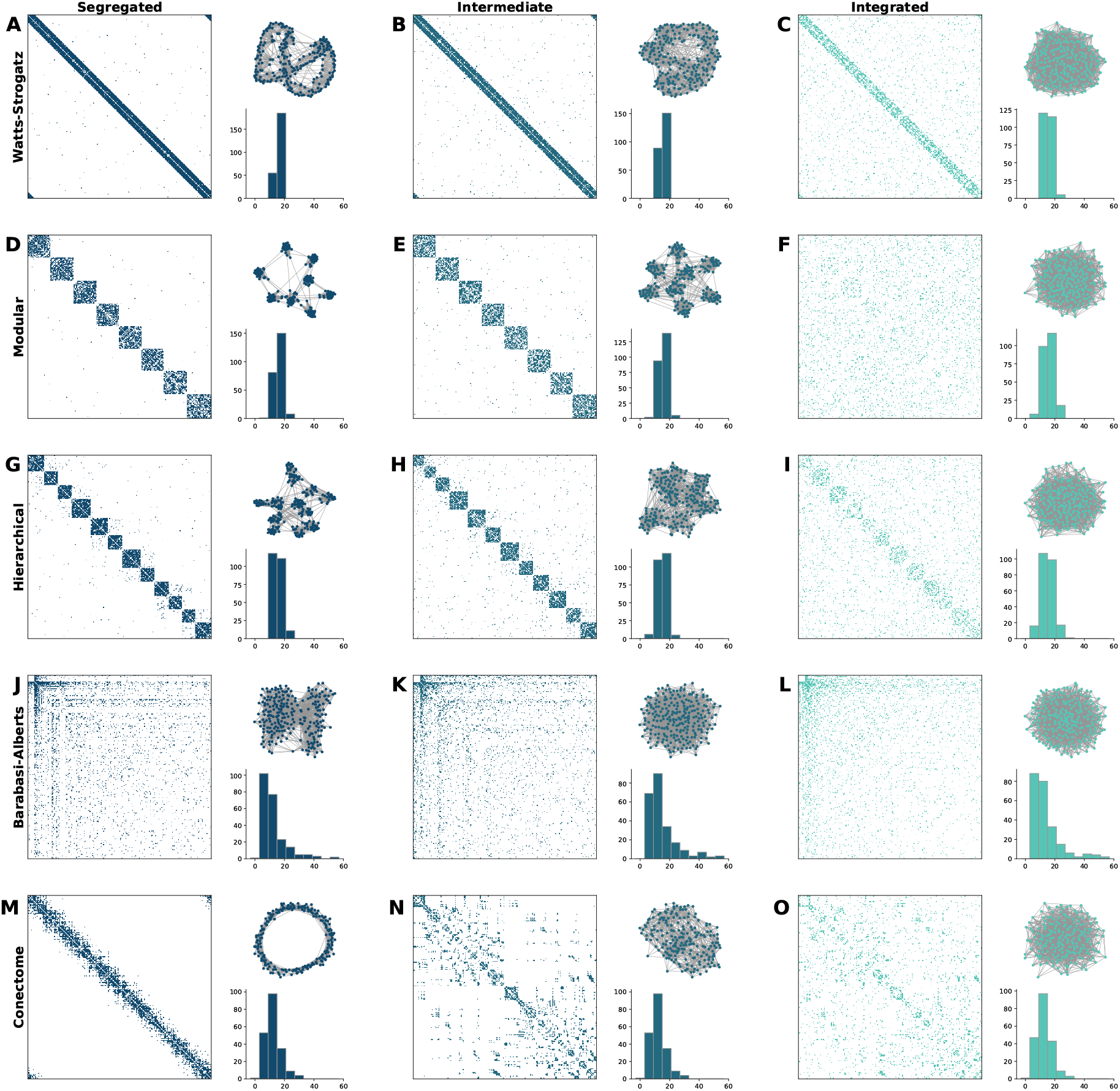
Network Types and Structural Properties: (A-C) Watts-Strogatz networks transition from highly ordered (segregated) to more randomized (integrated) structures by increasing the rewiring probability (*p*). (D-F) Modular networks show increasing inter-module connections, enhancing integration. (G-I) Hierarchical networks are constructed by iteratively adding inter-module connections in an ordered fashion to create a hierarchical structure. (J-L) Power-Law networks, generated with a modified Barabasi-Alberts algorithm, display variations in clustering by forming triangles. (M-O) Human connectome networks are modified to increase either segregation or integration using a custom algorithm. The networks span a spectrum from highly segregated to highly integrated, characterized by the Small-world index (*ω*).

### Integration/segregation is better described by *ω*

To characterize the degree of integration or segregation a network has, several metrics were calculated. Briefly, a segregated network is a network with a high amount of different communities. On the contrary, an integrated network could be characterized as a big single community.

Thus, Segregation (Clustering coefficient, modularity, and path length), and integration (global efficiency), metrics were calculated^24^, by using the brain connectivity toolbox for Python *bctp* ^1^

Additionally, a segregated (integrated) network is similar to a latticed (random) one^25^. As such, Telesford’s Small-World metric^26^, *ω*, was also calculated.

As our networks span a broad range in the Integration-segregation axis (Fig. 2 A), we seek to evaluate if a metric that discriminates between integration and segregation, that is *ω*, could establish the degree of the network’s integration/segregation.

**Figure 2.**
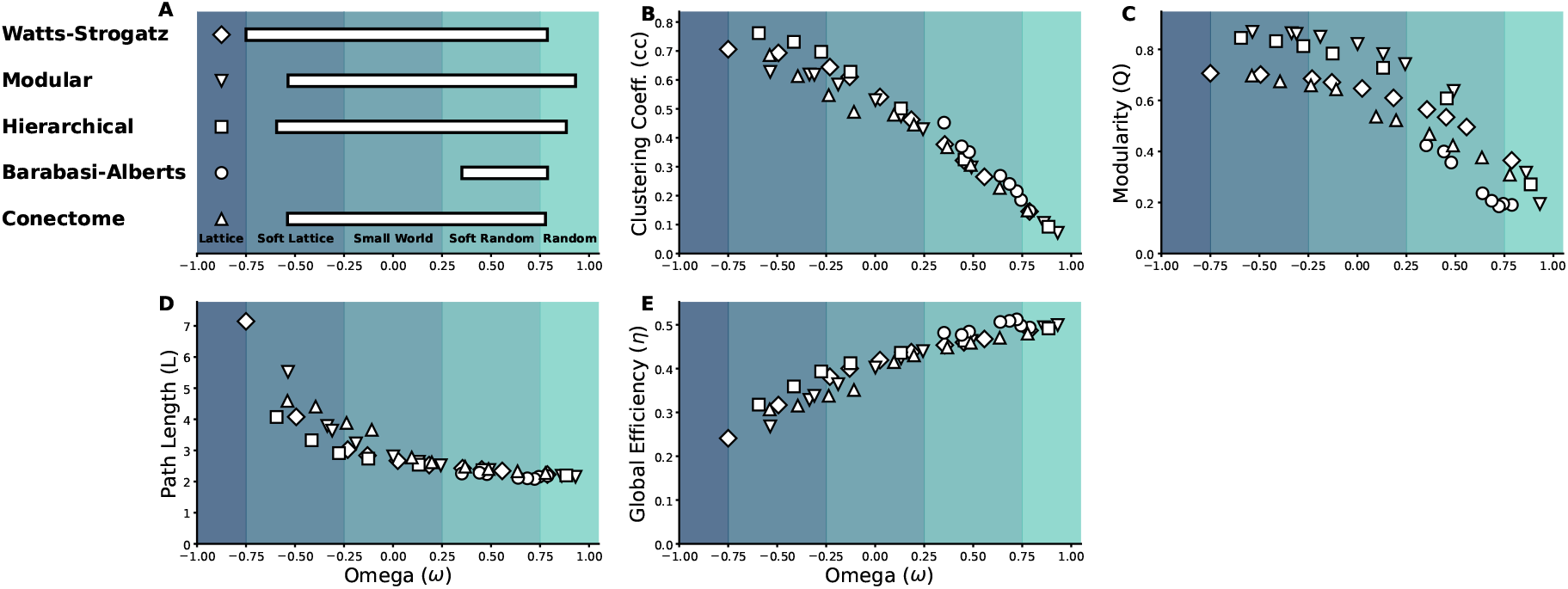
Integration and Segregation Metrics for different network topologies. (A) Network classification based on the Small-world index (*ω*) into lattice, soft lattice, small world, soft random, and random categories. (B-D) Relationship of *ω* with clustering coefficient, modularity, and path length, respectively. (E) Relationship of *ω* with global efficiency. Lattice: *ω* ≤ −0.75); soft lattice: −0.75 *< ω* ≤ −0.25; small world: 0.25 *< ω* ≤ 0.25; soft random: 0.25<*ω* ≤ 0.75; random: 0.75 *< ω*.

For this, we plotted each metric against *ω* (Fig. 2 B-D). We observed that the clustering coefficient decreases as *ω* increases, indicating a transition from locally clustered to more random structures (Fig. 2 B). Modularity, a measure of global segregation, also decreases with increasing *ω* (Fig. 2 C). Path length, which indicates network integration, shows an inverse relationship with *ω* (Fig. 2 D). Finally, global efficiency, representing network integration, increases with *ω* (Fig. 2 E).

In summary, *ω* correlates inversely with clustering coefficient and path length^25,26^, while global efficiency is directly correlated with *ω*^27^, showing a direct link between *ω* and the integration-segregation properties of the network.

### FCD is a proxy for network’s dynamical richness

To study how network topology determines the its dynamics, we used a Wilson-Cowan neural mass model^28^ on each node. The model considers a homeostatic plasticity mechanism that allows a better exploration of the parameter space, as the nodes maintain their oscillatory behavior within a wide range of external inputs. The dynamics of all networks were characterized at different values of the coupling strength *g*, distributed logarithmically between 0.01 and 2.5. Simulated signals were filtered, and downsampled. Hilbert Transform was used to obtain the phase and envelope in the 5-15 Hz band (figure 3 A). Time-resolved Functional Connectivity (FC) matrices were calculated as the pair-wise correlation between envelopes^29^, using a sliding window approach (window size = 2000 points, equivalent to 2 s; overlap = 75%; figure 3 A, bottom). Finally, the FCD matrix was calculated by computing the Euclidean distance between vectorized FCs (figure 3 B).

**Figure 3.**
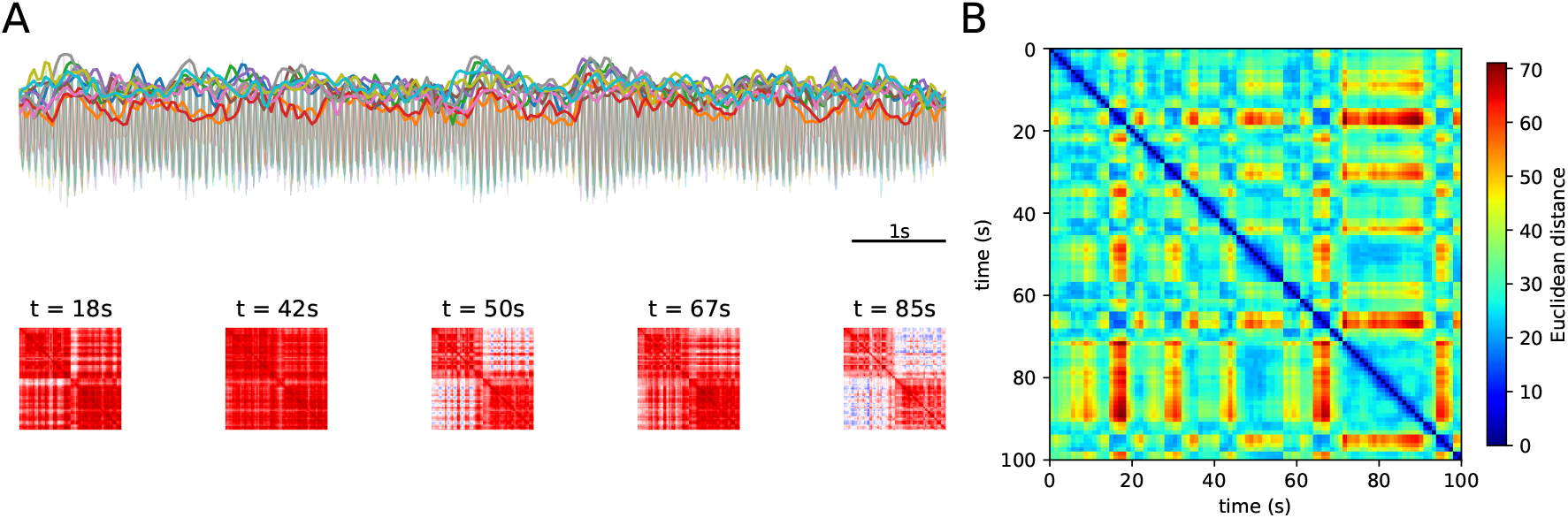
Simulation Workflow. **A** Trace signals(gray) and signal envelopes are shown. In red, FC networks for different times are shown. **b** To represent network dynamics, Functional Connectivity Dynamics was built by calculating the euclidean distance between vectorized FCs. Notice that *Var*(*FCD*_*g*=0.00*vg*≥0.1_) ≠ 0, however at intermediate values of g, *Var*(*FCD*) ≠ 0. Adapted from^16^

The FCD matrix shown in figure 3 B represents how different (or similar) are the FCs through which the network transits. Therefore, the FCD is a representation of the network dynamics^11^. When the coupling between nodes is *g* ≤ 0.04, the FCD appears as almost uniform green (figure 4 top), meaning that the difference between FCs is higher than 0 and mostly constant. Thus, the network is always found in different (un)synchronized states that are never revisited. On the other hand, when coupling *g* ≥ 1.0, the FCD matrix appears blue, meaning that the difference between FCs is zero and the synchronization pattern is only one static configuration throughout the simulation. Intermediate values of *g* cause the appearance of yellow and red patches in the FCD, meaning richer dynamics where FCs with both larger and smaller differences are observed (figure 4, top).

**Figure 4.**
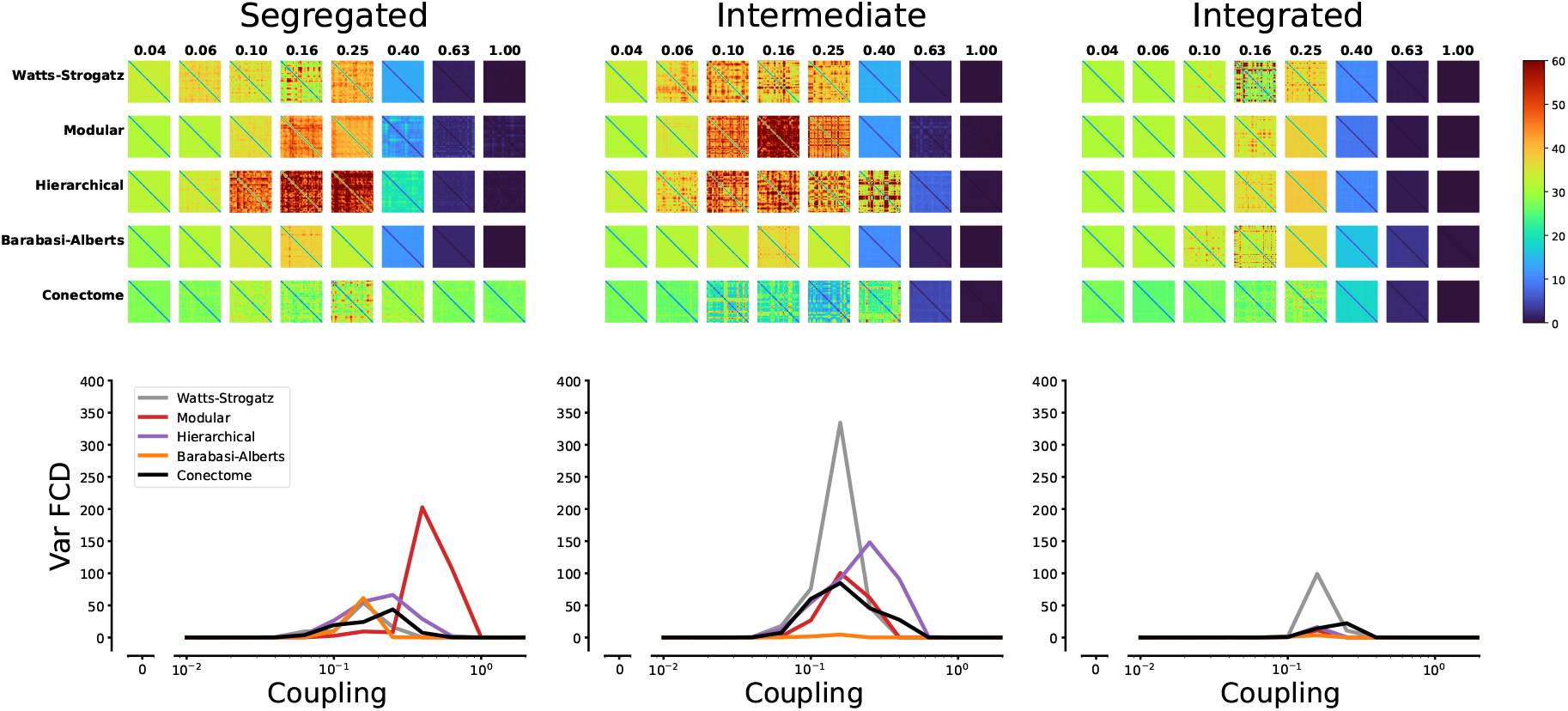
Network Topology drives Network Dynamics. The FCD patterns (top) of segregated (left), intermediate (middle), or integrated (right) networks are shown. Coupling value is shown to 1.00. Overall, intermediate networks are more dynamic, as their FC distance is higher (red patches). On the contrary, integrated networks exhibit less dynamics (green to blue patches). The variance of FCD is shown in the bottom plots. The X-axis represents coupling strength while the y-axis is the mean variance of FCD over 4 network realizations. Notice that Watts-Strogatz networks yield the higher values for Var(FCD) for intermediate or high levels of integration.

To account for the observed synchronization patterns, we use the variance of the off-diagonal values in the FCD, var(FCD). Var(FCD) is minimal at small or large *g* (indicating constant or fixed correlation patterns) and maximal when the FCD matrix has a patchy structure (Figure 4, bottom).

### Network Topology drives Network Dynamics

To account for the dynamical repertoire exhibited by each type of network, in the Integration/segregation continuum, we calculated the Var(FCD) for each global coupling value, and averaged the results of 10 different random seeds that governed the heterogeneity of the network for each network. Figure 4 depicts the FCD matrices for three characteristic networks of each type, and at different values of *g*. At the bottom, the line plots summarize the Var(FCD) over the *g* range explored. All networks showed a maximum dynamical richness at intermediate values of global coupling, as it has been previously established in simulation studies^16,20^. Also, networks classified as ‘intermediate’ between segregation and integration, show rich dynamics in a wider range of coupling values. Moreover, the networks of the Modular and Hierarchical types tend to show greater dynamics of FC.

### Network topology imposes Dynamical richness

To further explore the dynamical variations due to SC, we analyzed other two parameters that characterize network’s dynamic: Synchrony and Metastability (*χ*). Briefly, Synchrony measures the phase synchrony of all the signals of the network, ranging from no synchronization (0) to fully synchronized (1). Metastability (*χ*), on the other hand, measures the variability in time of the global synchrony^30^. Both measures are plotted along with Var(FCD) for all networks studied and across the whole range of global coupling in Figure 5. In all cases, there is a strong correspondence of shallow –sometimes staggered– synchronization curves with a higher occurrence of Metastability and Var(FCD). On the contrary, steep synchronization curves (typical for random networks) correlate with very small values of the measures associated with dynamical richness. Metastability and Var(FCD) mostly coincide for all networks but with some differences, such as in Connectome-based networks where the measures peak at clearly different values of coupling.

**Figure 5.**
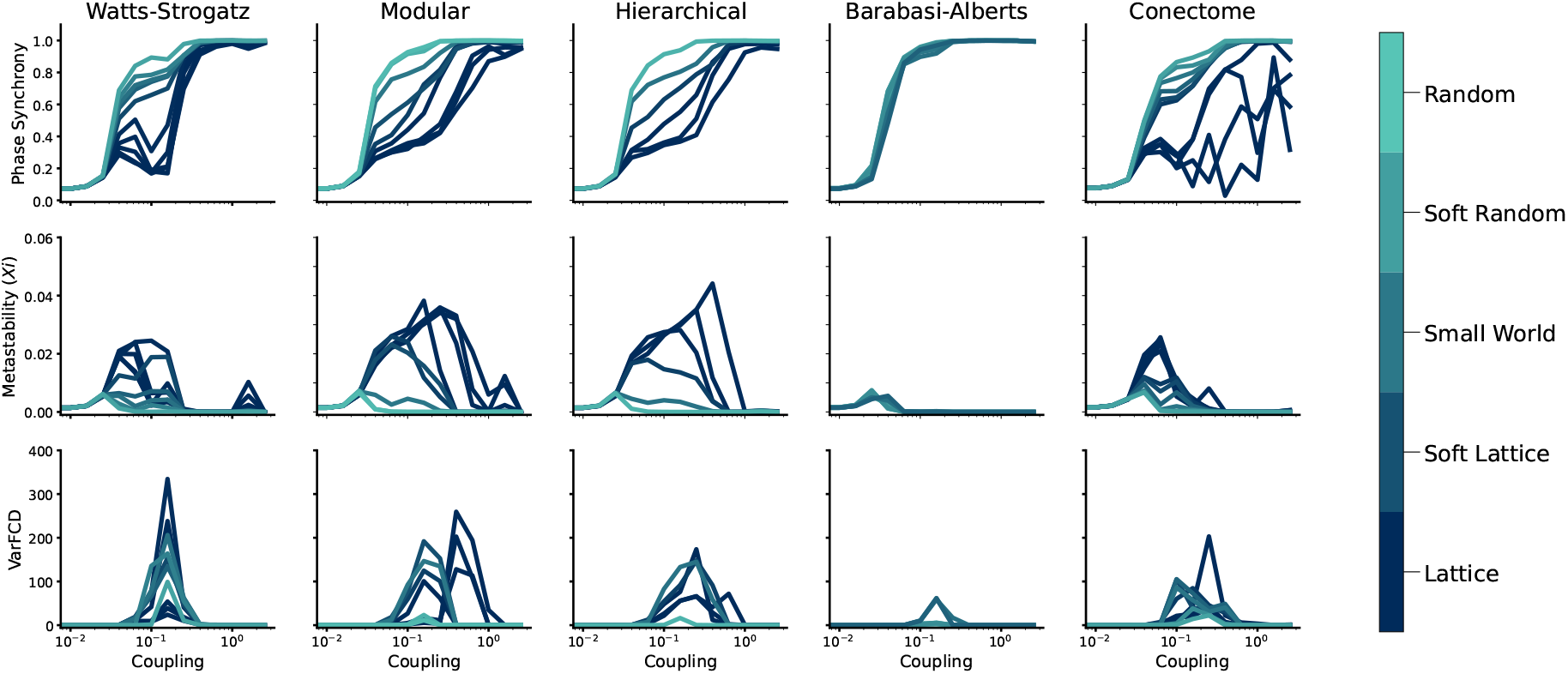
Network topology modulates Dynamical parameters. Network’s behavior for four dynamical metrics(rows), and different types of networks densities are shown. Colors represent different integration-segregation values ranging from latticed to random (*p* = 1.00). All values correspond to mean values over ten realizations.

### Linking structural features to Network Dynamics

For establishing a relationship between structure and dynamics, we reduced the dynamic range of each network (as shown in Figure 5) to a single value by calculating the area under the curve. This approach allowed the dynamics of any network, across the studied range of global coupling (*g*), to be summarized by a single parameter. Figure 6 illustrates the relationship between structural parameters and dynamical measures such as Metastability and Var(FCD).

**Figure 6.**
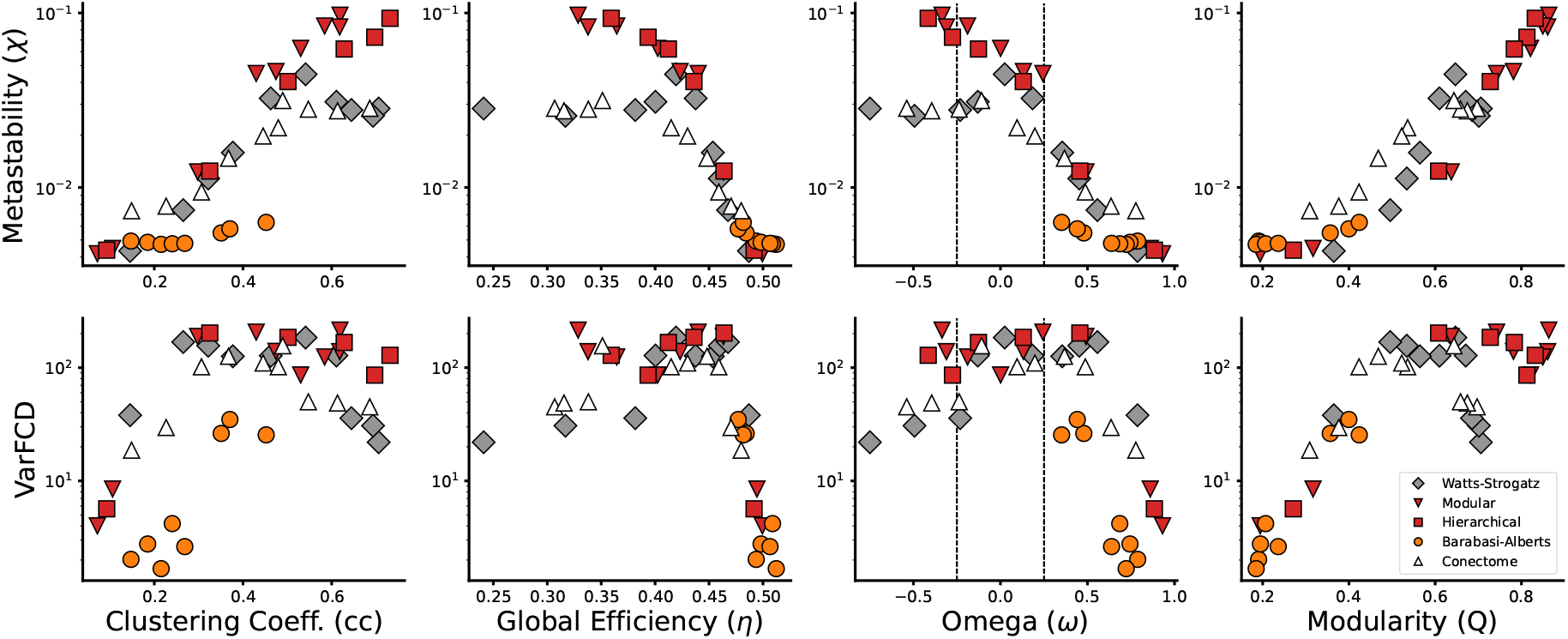
Relationship Between Structural Features and Network Dynamics. Metastability (*X*) and Var(FCD) plotted against clustering coefficient, global efficiency, and modularity for all network tested. Symbol shapes and colors denote the different types of networks, as shown in the legend. Vertical lines in the Omega plot denote the range considered as small-world (−0.25;0.25). Networks with small-world characteristics (*ω* close to zero) exhibit the highest dynamical richness, with Var(FCD) peaking at intermediate global coupling values. The plots highlight that structural features such as clustering coefficient, global efficiency, and modularity are key predictors of the dynamic behavior of networks, with small-world networks showing the optimal balance for dynamic flexibility.

The segregation measures tested—Clustering Coefficient (CC) (Figure 6, top left) and Modularity (Q) (Figure 6, top right)—both followed a monotonous exponential relationship with respect to *ξ*. However, the Clustering Coefficient (CC) showed distinct behavior in Barabási-Albert (scale-free) networks compared to other network types. At higher CC values, modular and hierarchical networks displayed trends that differed from those of Watts-Strogatz and connectome-based networks. Similarly, the integration metric (Global Efficiency, *η*) and the small-worldness index (*ω*) (Figure 6, middle-top) showed differing patterns for modular and hierarchical networks relative to Watts-Strogatz and connectome networks. This similarity between modular and hierarchical networks is unsurprising, as hierarchical networks are inherently modular in their structure.

On the other hand, Var(FCD) exhibited a broad inverted “U”-shaped relationship with structural features (Figure 6, bottom), peaking when the small-worldness index (*ω*) was between 0 and 0.5, indicating an optimal balance between integration and segregation. Var(FCD) declined sharply when integration increased (Global Efficiency) or segregation decreased (Modularity or Clustering Coefficient), emphasizing the need for segregated subnetworks to sustain dynamic synchronization patterns. Modularity (Q), a global segregation measure, better captured the increasing trend in Var(FCD) compared to the local Clustering Coefficient (CC), which showed deviations, particularly for Barabási-Albert networks. Notably, Watts-Strogatz and connectome networks experienced a significant drop in Var(FCD) at high segregation levels, unlike modular and hierarchical networks, which maintained high values and did not exhibit the inverted “U” trend.

### Modularity is a predictor for network Dynamics

Finally, to link structural parameters with dynamical properties, we calculated the mutual information (MI) between each structural parameter and the corresponding dynamical measures (Table 1 and Figure 7). Briefly, MI quantifies the relationship between two variables, *X* and *Y*, by measuring the amount of information about *Y* that can be gained by observing *X* ^31^. This method provided a model-agnostic framework to identify the structural parameter that best predicts the associated dynamical properties.

**Table 1.**
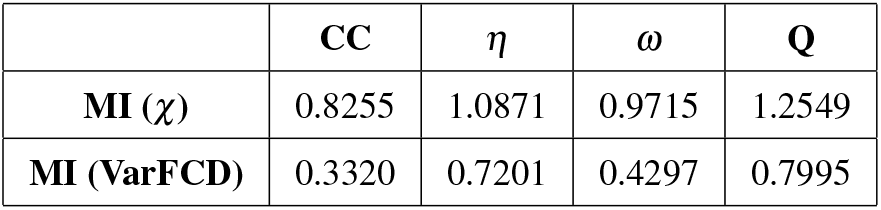
Mutual information values[nats] for structural and dynamical metrics.

**Figure 7.**
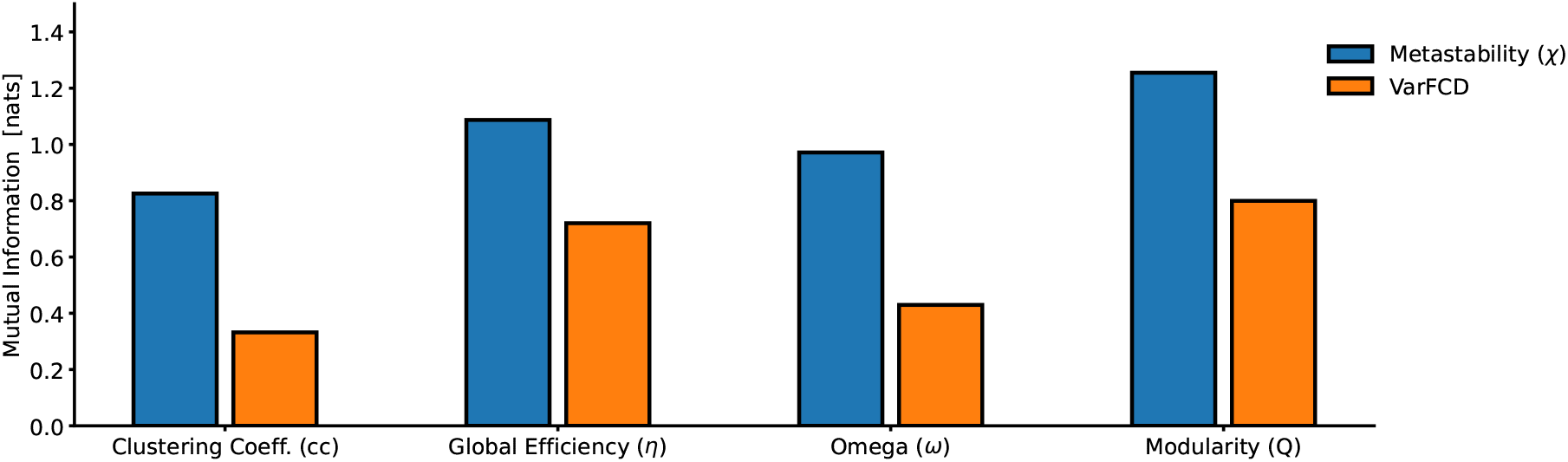
Mutual Information links network’s structure and dynamics. Bar shows the mutual information that each structural metric shares with dynamical ones, for all types of networks. Modularity (Q) is the structural parameter that shares more information with both dynamical parameter.

The mutual information (MI) values reveal distinct relationships between structural metrics and dynamic parameters. Metastability (*ξ*) consistently shows higher MI values across all metrics compared to Var(FCD), indicating that it is more strongly and predictably influenced by structural features. On the other hand, Var(FCD), is most effectively predicted by Efficiency and Modularity, highlighting their critical role in dynamic richness.

Among the structural metrics, Modularity (Q) exhibits the highest MI values for both *χ* and Var(FCD). This shows modularity’s role in mediating transitions across dynamical states by balancing localized specialization and global integration. Additionally, the relatively lower MI for *ω* in Var(FCD) compared to *χ* indicates that while small-worldness is significant for metastable states, it plays a less prominent role in dynamic functional connectivity variability, suggesting that modularity provides a more direct representation of the large-scale structural organization required to sustain rich and complex network dynamics.

Overall, global measures like Modularity (Q) and Efficiency (*η*) emerge as the most informative metrics for understanding network dynamics, particularly for the more complex patterns captured by Var(FCD).

## Discussion

In this study, we explored how network structure influences brain dynamics, focusing on metrics that quantify the integration-segregation balance, particularly modularity and small-worldness. We found that networks with an *ω* index close to zero or slightly positive –indicating small-world characteristics– exhibited the most dynamic behavior, as quantified by the variance of the FCD matrix. The networks that deviate from this tendency were found to have a high global efficiency, i.e. to be highly integrated. Additionally, the ability to transit from one dynamic state to another one was characterized by metastability *χ*^30^. Our experiments showed that highly segregated networks are more prone to transit to different states than integrated ones.

Previous research has extensively investigated the relationship between network structure and dynamics across different scales. At the local scale, random neural fluctuations and chaotic activity have been linked to phenomena such as dynamical wandering, where the brain transitions between various stable states over time^14–16,18,32^. At the mesoscale, mechanisms such as coupling between cortical areas, subnetworks, and delays between nodes have been proposed to drive state transitions by modifying the brain’s intrinsic, likely chaotic, oscillatory regime^19–21^. These studies collectively highlight the influence of structural connectivity on neural dynamics.

Building on these findings, our study focuses on the global scale, demonstrating that the presence of modules within a network, along with their interconnectivity, plays a crucial role in modulating global dynamics. By using mutual information to quantify the relationship between structural and dynamical parameters, we identified that structural features such as modularity is key for understanding how network structure shapes dynamic richness in a model-agnostic manner.

The relationship between structure and dynamics becomes evident when considering the level of modularity and interconnectivity in a network. At one extreme, networks composed of entirely disconnected modules exhibit flat dynamics, as limited communication between components leads to each module operating independently, with average distances between components too large to enable meaningful interaction. Conversely, enhancing communication between modules, for instance by reducing the mean path length, increases dynamical richness, enabling more complex interactions across the network. However, when interconnectivity becomes excessive, or modularity is absent as in random networks, the system transitions into a fully integrated state where all components synchronize, once again resulting in flat, homogeneous dynamics.

In addition to influencing dynamic richness, the presence of modules strongly correlates with high metastability. Networks with high modularity, such as modular or hierarchical networks, exhibit increased metastability, as the presence of distinct yet interacting modules creates conditions conducive to metastable regimes. This observation aligns with prior research demonstrating that modular organization promotes metastable dynamics^33^. Together, these findings underscore the critical role of modular structure in shaping both the richness and stability of network dynamics.

Our study also highlights the critical role of small-worldness in supporting the brain’s dynamical richness—its capacity to exhibit a wide range of dynamic states. The human connectome combines diverse topological characteristics, including small-world, modular, hierarchical, and power-law properties^34–37^. Among these, small-worldness stands out as particularly crucial because it balances integration (efficient global communication) and segregation (specialized local processing)^38^. Small-world networks, with their hallmark features of short path lengths and high clustering, uniquely facilitate both integration and segregation. Short path lengths enable rapid communication between distant brain regions, ensuring efficient global coordination, while high clustering supports local specialization within modules^34,39^. This dual capability surpasses the limitations of purely modular networks, which excel in segregation but lack efficient integration^40^, and powerlaw networks, which promote integration through highly connected hubs but lack strong local clustering. By combining these strengths, small-world networks provide the structural foundation for the brain’s dynamic flexibility and resilience.

Our analysis using Mutual Information (MI) shows that modularity is a stronger predictor of network dynamics compared to other structural metrics, such as small-worldness (*ω*). While *ω* is effective in representing the integration-segregation balance at a local level, modularity provides a clearer representation of the overall network architecture. Modularity describes the organization of distinct yet interconnected modules, which are closely tied to large-scale structural features that influence dynamic interactions. This characteristic makes modularity better suited to explain complex dynamic behaviors, including metastability (*ξ*) and dynamic functional connectivity variability (VarFCD). In contrast, the small-worldness index (*ω*), derived from the clustering coefficient (CC), emphasizes immediate node-level connections. Consequently, any limitations in MI observed for CC also affect *ω*, given their inherent connection. These findings suggest that modularity plays a key role in determining the dynamic properties of brain networks, offering insights that go beyond the capabilities of simpler metrics like clustering coefficient or global efficiency. The analysis illustrates how modular organization contributes to the complexity and adaptability of brain dynamics.

However, while these findings are promising, they should be interpreted with caution. One key limitation of our study is the exclusion of weighted networks, which restricts the generalizability of our conclusions. Prior research has shown that variations in coupling strength between brain regions significantly influence overall dynamics, particularly in individuals with Alzheimer’s disease^41^, major depressive disorder (MDD)^42^, cognitive decline^43^, or as part of the aging process^44^. Future studies should investigate the role of coupling strength in shaping neural dynamics, particularly at the macroscale level.

Additionally, while the variance of functional connectivity dynamics (VarFCD) provides valuable insights into how brain region connectivity evolves over time, it may not fully capture the complexity of brain dynamics. To achieve a more comprehensive understanding of dynamical richness, it is essential to explore higher-order interactions. These interactions, which account for simultaneous relationships among multiple brain regions, can uncover intricate patterns of connectivity and coordination that are otherwise overlooked^45^.

In conclusion, our findings suggest that highly modular networks are particularly adept at transitioning between distinct dynamical states, underscoring their crucial role in system segregation. The degree of interconnectivity between modules significantly shapes how networks navigate shifts between low and high activity levels. Notably, small-world networks exhibited the most dynamic behavior, as evidenced by the variance of the Functional Connectivity Dynamics (FCD) matrix.

While these results highlight the importance of structural metrics in capturing rich dynamical behavior, modularity emerged as the strongest predictor of network dynamics. Utilizing Mutual Information to assess these relationships provides a robust approach for identifying the most relevant structural parameters for predicting network dynamics, effectively capturing their interdependencies. The higher Mutual Information values associated with modularity, compared to other metrics, further reinforce its critical role in the mechanisms underlying dynamic richness and metastability

Given the study’s focus on binarized networks and pairwise correlations, future research could extend these findings by investigating weighted networks, where variations in connection strength may provide a deeper understanding of how modularity influences dynamic behavior. Additionally, incorporating higher-order interactions could offer deeper insights into the complex mechanisms underlying brain dynamics, capturing dependencies beyond pairwise relationships and enriching our understanding of network function.

## Methods

### Networks

Networks of 240 nodes (200 in human connectome) were created as such that each network has a density of 0.075. This means an average degree of 18 connections per node in the 240-node networks and 15 connections in the 200-node networks. All networks are binary and undirected, that is, the adjacency matrices are symmetric.

#### Watts-Strogatz Networks

These networks were generated using the Watts-Strogatz algorithm^25^. The algorithm starts with 240 nodes arranged in a circular lattice and each node is connected to its 18 nearest neighbors. Then, with probability *p*_*r*_, each connection is rewired to connect a randomly selected node within the network. The reconnection probability was varied from *p*_*r*_ = 0 (highly segregated lattice network) to *p*_*r*_ = 0.5 (highly integrated random network).

#### Modular Networks

This type of network consists of a set of internally topologically independent subnetworks (modules), which are connected to each other through a reduced number of (intermodule) links. 240 nodes were arranged in 8 modules of 40 nodes. Nodes within the modules were connected randomly with a density of 0.6. Then, with probability *p*_*inter*_, intra-module connections were replaced by inter-module connections, such that the degree of the initial modular network remains constant. The inter-module connection was varied between *p*_*inter*_ = 5 × 10^−4^ (highly segregated) to *p*_*inter*_ = 0.07 (highly integrated).

#### Hierarchical Networks

These networks where initially set up in a similar manner as Modular Networks. 240 nodes were divided in 12 modules and the nodes within the modules were randomly connected with density 0.9. In this case, the modules were of variable size between 16 and 24. Then, the hierarchical structure is implemented by iteratively connecting module pairs, with a probability that decreased as the hierarchy increased. Finally, a number of random connections were added (replacing existing ones to preserve the degree), to gradually increase the network integration.

#### Scale-free Networks

These networks where generated using the Holme and Kim algorithm^46^, as implemented in the NetworkX Python package^47^. This algorithm generates graphs with power-law degree distribution and a approximate average clustering coefficient. The algorithm begins with a small number of nodes (typically three) and grows the network by adding one node at a time. Each new node is connected to existing nodes with a preference for those that have higher degrees (rich-gets-richer). Additionally, after choosing a node based on its degree, the algorithm might create a triangle (triad closure) to increase clustering. In this way, by progressively increasing or decreasing the average clustering coefficient we obtained more segregated or integrated networks, respectively.

#### Human Connectome Networks

The Enigma toolbox^48^ was used to obtain a Human Connectome Project (HCP) connectome parcellated according to the Schaefer 200 parcellation. The weighted connectome was binarized using a threshold that left 7.5% of connections. To integrate the human connectome network, we use the **randmio_und_connected** function of the *Brain Connectivity Toolbox* (BCT) package implemented in Python (https://pypi.org/project/bctpy/). Starting by determining the number of rewiring iterations, we applied the function iteratively, increasing the rewires each time to move the network towards a randomized state where the integration is enhanced. The process is stopped once a desired integration level or a set number of iterations is reached, ensuring that while the network becomes more integrated, its degree distribution remains consistent.

To segregate the human connectome network, we developed an algorithm based on node modularity. It starts by computing an agreement matrix, derives a consensus partition for the network, and then calculates the nodal participation coefficient and z-scored modularity for each node. The core operation iterates over nodes, prioritizing those with high modularity scores. For these nodes, it identifies and deletes certain intra-module connections (those within the same community) and establishes new inter-module connections (those outside its community). This rewiring aims to change up to three connections per iteration, preserving the graph’s structure while emphasizing nodes with the highest modularity values.

### Structural Metrics

Integration and segregation in the generated networks were quantified by several metrics. *Global efficiency* is an estimator for network integration, being the average of the inverse of the shortest path lengths^49^. *Clustering coefficient* is a local measure of segregation, effectively counting how many triangles are formed when two neighbors of a node are connected as well^50^. *Modularity* measures segregation more globally, relying on the initial detection of modules and then measuring the balance between intra-module and inter-module links^51^. Finally, *small-world index ω* measures the integration/segregation balance by contrasting the Global Efficiency and Clustering Coefficient of each network with that of its corresponding latticed or random version^25^.

### Dynamical Model

#### Neural Mass Model: Wilson-Cowan with Plasticity

The cerebral dynamics of each constituent node in the obtained SC networks were simulated using an oscillatory neural mass model, the Wilson-Cowan model^52^, with the incorporation of an inhibitory synaptic plasticity (ISP) mechanism^28^. Unlike biophysically detailed models, this model provides a description of the evolution of average oscillatory, excitatory, and inhibitory activity within a neural mass (brain region). It represents a node in a large-scale neuronal network connected synaptically. This model employs only two differential equations, with key parameters being the strength of connectivity between each subtype of population (excitatory and inhibitory) and the input strength to each subpopulation. Varying these parameters generates diverse dynamic behaviors representative of brain activity, such as multistability, oscillations, traveling waves, and spatial patterns.

The activity in neural populations is governed by the equations^28^

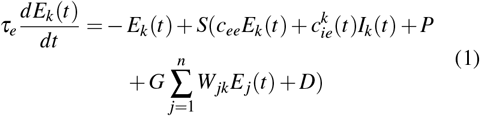

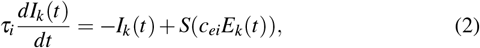

where *E*_*k*_ and *I*_*k*_ correspond to the average firing rates in excitatory and inhibitory populations in the *k*-th brain region (node in the network), respectively. *τ*_*e*_ and *τ*_*i*_ are the excitatory and inhibitory time constants, *c*_*ab*_ is the local connection strength from population *a* to population *b*, and *P* is the excitatory input constant. Long-range connections *W*_*jk*_ = {0, 1} from region *j* to region *k* are multiplied by a global coupling constant *G*. The constant *D* corresponds to additive noise, which is given by random values from a normal (Gaussian) distribution. Indices *k* and *j* run across the total number of nodes *n*. The nonlinear response function *S* is a sigmoid function given by:

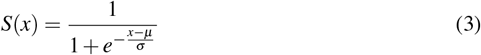

where *μ* and *σ* represent the threshold for the average firing rate and the variation in the threshold for neurons in each population, respectively.

The ISP mechanism is modeled as a time-dependent spike process, depending on both pre- and post-synaptic activity. The strength of the local inhibitory connection changes depending on the activity of the corresponding local excitatory population, according to

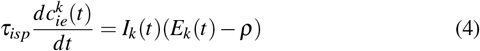

where *τ*_*isp*_ is the learning rate, and *ρ* is the target excitatory activity level, with initial value 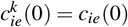. Without ISP, the local inhibitory coupling strength is constant and identical for all brain regions *k*, with 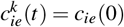.

#### Simulation

Based on the obtained sets of networks, the neuronal dynamics of each network were simulated using the Wilson-Cowan oscillatory neural mass model with plasticity in each node. The global coupling strength parameter *G* was varied over a logarithmic space range of values {0, ‥, 2.512}.

Simulations were performed for *t* = 102*s*, with a noisless transient of *t*_*trans*_ = 50*s* for ensuring system stabilization. Integration time step was *dt* = 0.0001. The first and last second of each simulation were discarded to remove mathematical artifacts that could influence subsequent stages.

To introduce heterogeneity in the obtained oscillatory signals among network nodes, the value of the excitatory input constant *P* was assigned randomly to each node, within a range of 0.3 − 0.5. The simulation parameters used were adjusted to ensure that the obtained signals exhibited oscillatory behavior. This simulation protocol was repeated 5 times (5 random seeds).

### Analysis

#### Signal preprocessing

Every simulated time serie was subsampled at 4 kHz, filtered with a lowpass filter for obtaining the subthreshold oscillation (Bessel 4th order, *ω*_*cut*_ =50Hz), downsampled to 200 Hz, and filtered again with a bandpass filter (Bessel, 4th order, *ω*_*cut*_ = *max* {*ω*_*i*_}± 3 Hz, where *w*_*i*_ is the mean frequency of HB oscillators). Hilbert transform was then applied for obtaining the instantaneous phase of the slow oscillation. The first and the last second of simulation were discarded in order to get rid of artifacts that Hilbert transform may generate. This signal was used for assesing dynamical analysis.

Functional connectivity was calculated as the pairwise correlation among all nodes.

### Dynamical Metrics

#### Functional Connectivity (FC)

The statistical dependency of neural signals was measured through a Functional Connectivity (FC) matrix^53^. Every element of this matrix corresponds to the pairwise phase synchrony between nodes *j, k*, averaged over a time window of duration *W* (*W*_*time*_ = 2*s*, overlap = 75 %).

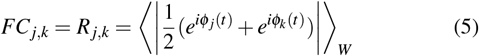

#### Functional Connectivity Dynamics (FCD)

Functional Connectivity Dynamics (FCD) matrix, which displays the dynamical repertoire of the system, was calculated using the Clarkson Distance between the vectorized lower triangular of *FC*_*t*_ and *FC*_*t*+*τ*_ as follows:

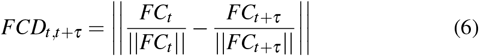

#### Synchrony

For assessing the overall synchrony of the network, kuramoto’s order parameter *R*(*t*) was calculated.

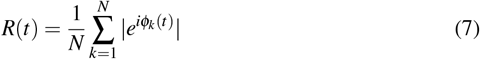

Where *i* is the complex unit and *ϕ*_*k*_(*t*) is the instantanous phase of the *k*-th node.

#### Metastability

Metastability (*χ*), understood as the switching between stable states^30^, was assessed by calculating the variance of *R*(*t*)

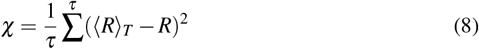

Where *τ* is the number of points that the signal *R* has, and ⟨*R*⟩_*T*_ is the average in time of the global synchrony.

#### Multistability

Multistability is the numerous stable points that a system has^30^. As such, we used the variance of FCD, VarFCD, as a proxy for quantifying multistability^16^.

### Mutual Information

The relationship between structural and dynamical parameters was evaluated using Mutual Information.

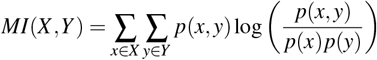

where, *p*(*x, y*) is the joint probability distribution, and *p*(*x*) and *p*(*y*) are the marginal distributions of *X* and *Y*, respectively.

For each structural parameter across the entire network set, Mutual Information with respect to multi- or metastability was computed using the scikit-learn function **mutual_info_regression** (https://scikit-learn.org/).

## Acknowledgements

J.PE. is recipient of a PhD Grant FIB-UV From UV. S.O.V. is recipient of a Ph.D. fellowship grant from ANID 21140603 (Chile). C.C. is supported by BrainLat/NIH and GBHI funding. P.O. is funded by the Advanced Center for Electrical and Electronic Engineering (AFB240002 ANID, Chile) and Fondecyt Regular Grant 1241469 (ANID, Chile).

## Author contributions statement

General Idea: JPE and PO.

Code and model creation: JPE, SOV, CCO, PO. Simulations: JPE, PO.

Analysis and plotting: JPE.

Writting, profreading, and manuscript approval: JPE,SOV,CCO, JPM, PO.

## Additional information

## Competing Interests

The authors declare no competing interests.

^1^https://pypi.org/project/bctpy/).

## References

1. Bullmore, E. & Sporns, O. The economy of brain network organization. Nat. Rev. Neurosci. 13, 336–349 (2012).

2. Kaiser, M. A tutorial in connectome analysis: topological and spatial features of brain networks. Neuroimage 57, 892–907 (2011).

3. Lynn, C. W. & Bassett, D. S. The physics of brain network structure, function and control. Nat. Rev. Phys. 1, 318– 332 (2019).

4. Sporns, O., Tononi, G. & Kötter, R. The human connectome: a structural description of the human brain. PLoS computational biology 1, e42 (2005).

5. Fornito, A., Zalesky, A. & Breakspear, M. The connectomics of brain disorders. Nat. Rev. Neurosci. 16, 159 (2015).

6. da Silva, F. L. Eeg and meg: relevance to neuroscience. Neuron 80, 1112–1128 (2013).

7. Van Den Heuvel, M. P. & Pol, H. E. H. Exploring the brain network: a review on resting-state fmri functional connectivity. Eur. neuropsychopharmacology 20, 519– 534 (2010).

8. Diez, I. et al. A novel brain partition highlights the modular skeleton shared by structure and function. Sci. reports 5, 10532 (2015).

9. Honey, C. J., Kötter, R., Breakspear, M. & Sporns, O. Network structure of cerebral cortex shapes functional connectivity on multiple time scales. Proc. Natl. Acad. Sci. 104, 10240–10245 (2007).

10. Honey, C. et al. Predicting human resting-state functional connectivity from structural connectivity. Proc. Natl. Acad. Sci. 106, 2035–2040 (2009).

11. Hansen, E. C., Battaglia, D., Spiegler, A., Deco, G. & Jirsa, V. K. Functional connectivity dynamics: modeling the switching behavior of the resting state. Neuroimage 105, 525–535 (2015).

12. Preti, M. G., Bolton, T. A. & Van De Ville, D. The dynamic functional connectome: State-of-the-art and perspectives. Neuroimage 160, 41–54 (2017).

13. Cabral, J., Kringelbach, M. L. & Deco, G. Functional connectivity dynamically evolves on multiple time-scales over a static structural connectome: Models and mechanisms. NeuroImage 160, 84–96, DOI: 10.1016/j.neuroimage.2017.03.045 (2017).

14. Berglund, N. & Gentz, B. Stochastic dynamic bifurcations and excitability. Stoch. methods Neurosci. 64–93 (2010).

15. Heitmann, S. & Breakspear, M. Putting the “dynamic” back into dynamic functional connectivity. Netw. Neurosci. 2, 150–174 (2018).

16. Orio, P. et al. Chaos versus noise as drivers of multistability in neural networks. Chaos: An Interdiscip. J. Nonlinear Sci. 28, 106321 (2018).

17. Piccinini, J. et al. Noise-driven multistability vs deterministic chaos in phenomenological semi-empirical models of whole-brain activity. Chaos (Woodbury, N.Y.) 31, 023127, DOI: 10.1063/5.0025543 (2021).

18. Xu, K., Maidana, J. P., Castro, S. & Orio, P. Synchronization transition in neuronal networks composed of chaotic or non-chaotic oscillators. Sci. reports 8, 8370 (2018).

19. Deco, G., Jirsa, V., McIntosh, A. R., Sporns, O. & Kötter, R. Key role of coupling, delay, and noise in resting brain fluctuations. Proc. Natl. Acad. Sci. 106, 10302–10307 (2009).

20. Deco, G. & Jirsa, V. K. Ongoing cortical activity at rest: criticality, multistability, and ghost attractors. J. Neurosci. 32, 3366–3375 (2012).

21. Cabral, J. et al. Exploring mechanisms of spontaneous functional connectivity in meg: how delayed network interactions lead to structured amplitude envelopes of band-pass filtered oscillations. Neuroimage 90, 423–435 (2014).

22. Batista-García-Ramó, K. & Fernández-Verdecia, C. I. What we know about the brain structure–function relationship. Behav. Sci. 8, 39 (2018).

23. Humphries, M. D. & Gurney, K. Network ‘small-worldness’: a quantitative method for determining canonical network equivalence. PloS one 3, e0002051 (2008).

24. Rubinov, M. & Sporns, O. Complex network measures of brain connectivity: uses and interpretations. Neuroimage 52, 1059–1069 (2010).

25. Watts, D. J. & Strogatz, S. H. Collective dynamics of ‘small-world’networks. nature 393, 440 (1998).

26. Telesford, Q. K., Joyce, K. E., Hayasaka, S., Burdette, J. H. & Laurienti, P. J. The ubiquity of small-world networks. Brain connectivity 1, 367–375 (2011).

27. Achard, S. & Bullmore, E. Efficiency and cost of economical brain functional networks. PLoS computational biology 3, e17, DOI: 10.1371/journal.pcbi.0030017 (2007).

28. Abeysuriya, R. G. et al. A biophysical model of dynamic balancing of excitation and inhibition in fast oscillatory large-scale networks. PLOS Comput. Biol. 14, e1006007, DOI: 10.1371/journal.pcbi.1006007 (2018).

29. Hipp, J. F., Hawellek, D. J., Corbetta, M., Siegel, M. & Engel, A. K. Large-scale cortical correlation structure of spontaneous oscillatory activity. Nat. neuroscience 15, 884–890 (2012).

30. Kelso, J. S. Multistability and metastability: understanding dynamic coordination in the brain. Philos. Transactions Royal Soc. B: Biol. Sci. 367, 906–918 (2012).

31. Timme, N. M. & Lapish, C. A tutorial for information theory in neuroscience. eneuro 5 (2018).

32. Piccinini, J. et al. Noise-driven multistability vs deterministic chaos in phenomenological semi-empirical models of whole-brain activity. Chaos: An Interdiscip. J. Nonlinear Sci. 31 (2021).

33. Hizanidis, J., Kouvaris, N. E., Zamora-López, G., Díaz-Guilera, A. & Antonopoulos, C. G. Chimera-like states in modular neural networks. Sci. reports 6, 19845 (2016).

34. Bassett, D. S. & Bullmore, E. T. Small-world brain networks revisited. The Neurosci. 23, 499–516 (2017).

35. Sporns, O. & Betzel, R. F. Modular brain networks. Annu. review psychology 67, 613–640 (2016).

36. Meunier, D., Lambiotte, R., Fornito, A., Ersche, K. & Bullmore, E. T. Hierarchical modularity in human brain functional networks. Front. neuroinformatics 3, 571 (2009).

37. Tomasi, D. G., Shokri-Kojori, E. & Volkow, N. D. Brain network dynamics adhere to a power law. Front. Neurosci. 11, 72 (2017).

38. Wang, R. et al. Segregation, integration, and balance of large-scale resting brain networks configure different cognitive abilities. Proc. Natl. Acad. Sci. 118, e2022288118 (2021).

39. Bassett, D. S. & Bullmore, E. Small-world brain networks. The neuroscientist 12, 512–523 (2006).

40. Cohen, J. R. & D’Esposito, M. The segregation and integration of distinct brain networks and their relationship to cognition. J. Neurosci. 36, 12083–12094 (2016).

41. Schumacher, J. et al. Dynamic functional connectivity changes in dementia with lewy bodies and alzheimer’s disease. NeuroImage: Clin. 101812 (2019).

42. Javaheripour, N. et al. Altered brain dynamic in major depressive disorder: state and trait features. Transl. Psychiatry 13, 261 (2023).

43. Wei, Y.-C. et al. Functional connectivity dynamics altered of the resting brain in subjective cognitive decline. Front. Aging Neurosci. 14, 817137 (2022).

44. Jauny, G. et al. Linking structural and functional changes during aging using multilayer brain network analysis. Commun. Biol. 7, 239 (2024).

45. Gatica, M. et al. High-order interdependencies in the aging brain. Brain connectivity 11, 734–744 (2021).

46. Holme, P. & Kim, B. J. Growing scale-free networks with tunable clustering. Phys. Rev. E 65, 026107, DOI: 10.1103/PhysRevE.65.026107 (2002).

47. Hagberg, A. A., Schult, D. A. & Swart, P. J. Exploring network structure, dynamics, and function using networkx. In Varoquaux, G., Vaught, T. & Millman, J. (eds.) Proceedings of the 7th Python in Science Conference, 11 – 15 (Pasadena, CA USA, 2008).

48. Larivière, S. et al. The ENIGMA Toolbox: multiscale neural contextualization of multisite neuroimaging datasets. Nat. Methods 18, 698–700, DOI: 10.1038/s41592-021-01186-4 (2021).

49. Latora, V. & Marchiori, M. Efficient behavior of smallworld networks. Phys. review letters 87, 198701 (2001).

50. Onnela, J.-P., Saramäki, J., Kertész, J. & Kaski, K. Intensity and coherence of motifs in weighted complex networks. Phys. Rev. E 71, 065103 (2005).

51. Clauset, A., Newman, M. E. & Moore, C. Finding community structure in very large networks. Phys. review E 70, 066111 (2004).

52. Wilson, H. R. & Cowan, J. D. Excitatory and inhibitory interactions in localized populations of model neurons. Biophys. journal 12, 1–24 (1972).

53. Messé, A., Hütt, M.-T., König, P. & Hilgetag, C. C. A closer look at the apparent correlation of structural and functional connectivity in excitable neural networks. Sci. reports 5, 7870 (2015).

